# LOca-1: A minimal calcium-activated arginyl-specific orthocaspase from the Loki’s Castle hydrothermal vent field

**DOI:** 10.64898/2026.11.04.764195

**Authors:** Hasan Arsın, Petra Hribovšek, Liam Urich, Anita-Elin Fedøy, Runar Stokke, Paweł Mak, Anna-Karina Kaczorowska, Anthony J O’Donoghue, Tadeusz Kaczorowski, Ida Helene Steen

**Author notes:** Corresponding Authors IHS HA.

## Abstract

Orthocaspases are prokaryotic homologs of metacaspases within the C14 family of cysteine proteases, yet their biochemical properties and cellular roles remain poorly understood. Here, we report the discovery and comprehensive characterization of LOca-1, a metagenome-derived orthocaspase from the Loki’s Castle hydrothermal vent field. LOca-1 consists primarily of a single, conserved p20 catalytic domain and undergoes calcium-dependent activation through cleavage at a conserved arginine residue. Once activated, LOca-1 displays strict arginyl-specific proteolytic activity and strong selectivity for residues surrounding the cleavage site. The enzyme is both thermoactive and thermostable, exhibiting maximal activity at 65 °C and a melting temperature of approximately 72 °C. Structural modelling revealed conservation of the catalytic core relative to plant metacaspases despite substantial sequence divergence. To our knowledge, LOca-1 is only the second biochemically characterized orthocaspase and the first reported from a deep-sea hydrothermal vent environment. LOca-1 expands the known biochemical diversity of prokaryotic metacaspase homologs and provides insights into the properties of a minimal p20-domain orthocaspase. Together, our findings establish LOca-1 as a thermostable, calcium-activated arginyl-specific orthocaspase and provide a foundation for future studies of orthocaspase function in prokaryotes.

## Introduction

Within the broad proteolytic landscape, the C14 family of cysteine proteases represents an ancient, evolutionarily conserved group defined by the caspase/hemoglobinase fold and a catalytic His–Cys dyad [1]. Despite this shared structural core, C14 proteases show significant variation in activation mechanisms, substrate specificity, and biological function. Canonical caspases, restricted to metazoans, are aspartate-specific and mediate apoptosis and inflammatory signalling, whereas metacaspases, found in plants, fungi, protists, and some bacteria, prefer cleaving after basic residues (Arg/Lys) and participate in a range of non-apoptotic processes including stress responses and proteostasis [2,3]. Paracaspases, present in metazoans and slime moulds, also cleave after arginine but function primarily in signalling pathways rather than in cell death [4–6].

Recent comparative genomic studies have further revealed that caspase-fold proteases predate eukaryotes and are widespread in bacteria, leading to the identification of orthocaspases as the prokaryotic members of the C14 family [7]. Orthocaspases retain the conserved p20 catalytic domain but typically lack the canonical metacaspase p10 subunit. They frequently occur in various multidomain architectures that extend beyond the protease core, reflecting substantial structural diversification [5]. Large-scale studies have also shown that orthocaspases are unevenly distributed across bacterial lineages [8,9]. Interestingly, orthocaspases are frequently accompanied by pseudo-orthocaspases lacking the catalytic dyad. The biological significance of these homologs remains unresolved [10,11].

Despite their apparent prevalence and architectural diversity, experimental data on orthocaspases remain scarce. To date, only a single orthocaspase, MaOC1 from the freshwater cyanobacterium *Microcystis aeruginosa*, has been biochemically characterized. MaOC1 was shown to be a proteolytically active, arginyl-specific cysteine protease that undergoes calcium-independent activation via autocatalytic processing [7]. More recently, its biological function was linked to cleaving an antitoxin within a toxin–antitoxin system, supporting a regulatory proteolytic role [12]. However, in addition to the conserved p20 domain, MaOC1 also possesses an unresolved C-terminal extension [2,9]. Therefore, characterizing a minimal, single-domain enzyme remains essential for defining the intrinsic biochemical properties of the catalytic p20 core.

Deep-sea hydrothermal vent environments are a rich source of enzymes with adaptations to unusual and fluctuating physicochemical conditions, including steep gradients in temperature, redox potential, and ionic composition [13,14]. Proteins derived from these environments frequently exhibit enhanced intrinsic stability and other adaptations to extreme physicochemical conditions, making hydrothermal systems valuable for enzyme discovery and investigating fundamental biochemical mechanisms [15–21].

Here, we report the discovery and comprehensive biochemical characterization of LOca-1 (Loki’s Castle Orthocaspase-1), a metagenome-derived orthocaspase identified from the Loki’s Castle hydrothermal vent field along the Arctic Mid-Ocean Ridges [22]. By combining enzymatic assays, structural modelling, and phylogenetic analyses, we demonstrate that LOca-1 is a thermostable, calcium-activated, arginyl-specific orthocaspase with a minimal p20-domain architecture. These findings provide new insight into the intrinsic biochemical properties of the conserved p20 catalytic domain, which forms the functional core of all orthocaspases.

## Methods

### The source and phylogenetic placement of LOca-1 from Loki’s Castle vent field

The LOca-1 gene was identified from a metagenomic assembly (Chimney19) from a black smoker chimney at the Loki’s Castle hydrothermal vent field [22]. Due to its location on a short contig comprising only three predicted genes, no reliable taxonomic assignment could be made.

The enzyme was therefore designated LOca-1: Lokís Castle Orthocaspase 1 to conform to the suggested common nomenclature of orthocaspases [23]. Closest homologs of LOca-1 were identified by blastp alignment using two databases with defined thresholds. For the GenBank ClusteredNR environmental sequences database, hits were retained at ≥94% query cover, ≥coverage, 47% sequence identity. For the RefSeq protein database (refseq_protein), hits were retained at ≥94% query cover and ≥coverage40% sequence identity threshold (as of July 2025). These thresholds were chosen to capture divergent orthocaspase homologs while excluding unrelated C14 proteases.

Sequences were aligned using MAFFT L-INS-i v7.525 [24], trimmed with trimAl v1.5 (– gappyout) [25], and manually curated in AliView v1.30 [26]. Phylogenetic relationships were inferred using IQ-TREE v2.4.0 [27] based on an alignment of 186 sequences spanning 266 positions. Branch support was assessed using SH-like approximate likelihood ratio tests [28] and ultrafast bootstrapping [29] with 1000 replicates. The best-fit substitution model (Q.pfam+I+R5) was selected using ModelFinder [30]. The tree was rooted at the midpoint. Environmental and taxonomic information for each entry were obtained from the available metadata in NCBI[31].

### Sequence alignment of LOca-1 with reference metacaspases

A multiple amino acid sequence alignment was generated, including the full-length LOca-1 sequence (288 aa) and representative reference metacaspases. Reference sequences comprised the closest structural and functional homologs identified from the Protein Data Bank (PDB_mmCIF70_25_May) [32,33] and the MEROPS database (Release 12.5) [34], including metacaspase-9 from *Arabidopsis thaliana* (AtMCA9; UniProt ID Q9FYE1) and metacaspase-1 from *Ajellomyces capsulatus* (AcMca-1; UniProt ID A6R7B8). Sequences were aligned using MAFFT L-INS-i v7.525 [24](Katoh and Standley, 2013) with a gap-opening penalty of 0.9. The resulting alignment was visualized using ESPript 3.0 [35].

### Structural model of LOca-1 and comparison with the crystal structure of *AtMCA9*

The amino acid sequence of LOca-1 (S1 File) was submitted to the AlphaFold3 server for three-dimensional structure prediction [36]. The resulting model was compared with its closest structural homolog identified in the Protein Data Bank (PDB) using the HHpred pipeline [37], namely the crystal structure of *Arabidopsis thaliana* metacaspase 9 (AtMCA9; PDB ID: 8A53) [38]. Structural superposition and visualization were performed using ChimeraX [39].

### Recombinant protein production and purification

A codon-optimized synthetic gene coding for LOca-1 and a mutant carrying a catalytic cysteine substitution (C138A) were synthesized by Thermo Fisher Scientific (Vilnius, Lithuania) for heterologous expression in *Escherichia coli*. Both genes were cloned into the pET-303/CT-His expression vector using XbaI and XhoI restriction sites, and where alanine was added before the XhoI restriction site, yielding recombinant proteins containing a C-terminal hexahistidine tag (ALEHHHHHH).

The resulting expression plasmids were transformed into *E. coli* BL21-Gold(DE3) competent cells (Agilent Technologies, Waldbronn, Germany) using a standard heat-shock transformation protocol (Pope & Kent, 1996). Transformed cells were plated on LB agar supplemented with 100 µg mL⁻¹ ampicillin and incubated overnight at 37 °C. Single colonies were used to inoculate LB pre-cultures containing 100 µg mL⁻¹ ampicillin, which were grown for 20 h at 37 °C with shaking at 200 rpm. Expression cultures were initiated the following day using a 2.5% (v/v) inoculum. After 2 h of growth at 37 °C, cultures reached an OD₆₀₀ of approximately 0.65 and were cooled to 20 °C prior to induction with 0.5 mM IPTG (final concentration).

Cultures were then incubated for an additional 20 h. Cells were harvested by centrifugation at 6,000 × g for 15 min (Allegra™ 21R centrifuge, Beckman Coulter, Krefeld, Germany), and cell pellets were stored at −20 °C. Pellets were resuspended in lysis buffer containing 40 mM HEPES (pH 8.0), 250 mM NaCl, 0.5% (v/v) Tween-20, 4% (v/v) glycerol, and 15 mM imidazole, then incubated on ice for 15 min. Cells were lysed by ultrasonication, and insoluble material was removed by centrifugation at 4,000 × g for 10 min.

LOca-1 was purified from the soluble fraction by immobilized metal affinity chromatography (IMAC) using a HisTrap™ FF column operated on an ÄKTA Start system (Cytiva, Uppsala, Sweden). Buffer A consisted of 40 mM HEPES (pH 8.0), 250 mM NaCl, 0.5% (v/v) Tween-20, 4% (v/v) glycerol, and 15 mM imidazole, whereas Buffer B contained the same components supplemented with 500 mM imidazole. Elution fractions containing LOca-1, identified from the chromatographic profile, were pooled, and the purity of the pooled protein preparation was assessed by SDS–PAGE. The pooled fractions were then concentrated and subjected to buffer exchange using Amicon Ultra centrifugal filters (10 kDa molecular weight cutoff; Merck Millipore, Darmstadt, Germany) into a storage buffer containing 40 mM HEPES (pH 8.0), 250 mM NaCl, and 4% (v/v) glycerol.

### Activation of the LOca-1 zymogen and detection of peptidase activity

To exhibit proteolytic activity, LOca-1 must first undergo calcium-dependent activation. Both LOca-1 and the C138A variant were diluted in an activation buffer (50 mM HEPES pH 7.5, 75 mM NaCl, and 1 mM CaCl_2_) to a final concentration of 0.2 mg/mL. The enzymes were then incubated at 60 °C for 0, 2.5, 5 and 10 minutes, and immediately transferred to ice. To visualize the results, SDS-PAGE was performed using 15 µL of each sample (∼3 µg protein) with 5 µL of loading dye. The non-incubated samples (t=0) were mixed with activation buffer on ice and were immediately loaded onto the gel. The common thermal denaturation step was omitted to avoid autoproteolysis.

The proteolytic activity of LOca-1 was initially assayed using the arginyl-bond-containing fluorescent peptide substrate Boc-VPR-amc. This peptide substrate was chosen for its previously demonstrated effectiveness for assaying other arginine-targeting cysteine proteases, such as globupain [19,40]. 50 µL of 2.9 µg/mL LOca-1 was mixed with 50 µL of 100 µM peptide substrate, both prepared in activation buffer, to achieve final CaCl_2_ concentrations ranging from 0 to 20 mM. The resulting reaction mixtures had final concentrations of 1.45 µg/mL enzyme and 50 µM peptide substrate. A negative control, in which the enzyme was replaced with assay buffer, was used to measure baseline fluorescence. Each sample was run in triplicate. The assay was run in a black 96-well plate within an EnSpire™ 2300 Multilabel Reader (PerkinElmer, Turku, Finland), set to incubate at 60 °C, with fluorescence measured every 20 sec at excitation and emission wavelengths of 360 and 460 nm, respectively. After measurement, raw data were processed in GraphPad Prism (version 10.4.1) to remove the baseline, calculate errors, and create figures. Unless otherwise specified, all error values are presented as the standard error of the mean (SEM).

Furthermore, LOca-1 was assayed in the presence of several common divalent metal ions to determine whether activation could be induced by cations other than calcium. For these experiments, the CaCl₂ present in the activation buffer was replaced with 5 mM concentrations of MgCl₂, MnCl₂, ZnCl₂, CuCl₂, or CoCl₂. Enzyme activation and peptidase activity were subsequently monitored using the Boc-VPR-AMC fluorogenic substrate assay as described above.

### N-terminal sequencing

The N-terminal amino acid sequences of LOca-1 fragments generated upon activation were determined using a PPSQ31A automatic protein sequencer (Shimadzu, Kyoto, Japan). Protein fragments were separated by SDS-PAGE, electrotransferred onto a polyvinylidene difluoride (PVDF) membrane, and subjected to Edman degradation sequencing.

### LOca-1 C138A catalytic variant

A substitution variant of LOca-1 (C138A) was designed to disable the enzymés predicted active cysteine and to serve as a benchmark for clarifying its activation mechanism and peptidase activity. Three aliquots of the mutant enzyme were prepared in the same activation buffer at the concentrations used above. A negative control was prepared without calcium and kept on ice throughout the experiment. To assess trans activation, one of the two other aliquots was spiked with ∼0.8 µg of WT LOca-1. These samples contained 1 mM CaCl_2_ and were incubated at 60 °C for 5 min before SDS-PAGE analysis, which was performed as described above.

To confirm the absence of activity resulting from substitution of the active enzyme, proteolytic activity was assayed against the substrate Boc-VPR-amc alongside the wild-type enzyme. Both enzyme forms were activated in the activation buffer at 60 °C for 5 min, then diluted to ∼0.2 µg/mL in the same buffer, yielding a final concentration of ∼0.1 µg/mL in the reaction mix. The assay was carried out as described above, and the resulting fluorescence, after baseline removal, was plotted at 2000 s using GraphPad Prism.

### Thermal stability analysis via Nano DSF

The thermostability of 1 mg/mL LOca-1 zymogen was assessed using nanoscale differential scanning fluorimetry (nanoDSF) in a buffer containing 40 mM HEPES (pH 8.0), 250 mM NaCl, and 5% glycerol. Measurements were performed on the Prometheus NT.48 AGOHTU instrument (NanoTemper Technologies, Munich, Germany) using PR.ThermControl software and standard-grade capillaries. Prior to analysis, capillaries were sealed with sealing paste according to the manufacturer’s instructions. Protein unfolding was monitored over a temperature range of 20-110 °C at a heating rate of 1 °C/min. Unfolding transitions were detected by changes in the intrinsic fluorescence of tryptophan and tyrosine at emission wavelengths of 330 and 350 nm. Protein aggregation was monitored simultaneously by back- reflection optics, based on measuring changes in sample turbidity caused by thermally induced aggregation. Melting temperatures (*T*_m_), unfolding onset temperatures, and aggregation temperatures (Tagg) were determined using PR.ThermControl and PR.StabilityAnalysis software (NanoTemper Technologies). All measurements were performed in triplicate.

### Analytical ultracentrifugation

Sedimentation velocity experiments were performed in a Beckman-Coulter ProteomeLab XL- I analytical ultracentrifuge (Indianapolis, IN, United States), equipped with an AN 60Ti 4-hole rotor and 12 mm path-length, double-sector charcoal-Epon cells, loaded with 400 μL of samples and 410 μL of buffer (40 mM HEPES pH 8.0, 250 mM NaCl, and 5% glycerol). To stabilize the temperature, the rotor was held at 20 °C for 60 min before the run. The experiment was conducted at 20 °C and 50,000 rpm, with LOca-1 at a concentration corresponding to OD_280_=0.16, in continuous-scan mode with a radial spacing of 0.003 cm. 100 absorbance scans were collected at 280 nm at 4-minute intervals. The buffer’s biophysical parameters: density (1,02357 g/mL) and viscosity (0.01184 Poise) were measured at 20 °C using an Anton Paar DMA 5000 density meter and a Lovis 2000 ME viscometer. Protein partial specific volume (V- bars) was estimated at 0.7330 mL/g using SEDNTERP software (version 1.10, Informer Technologies Inc., Dallas, TX, United States). Data were analysed using the “Continuous c(s) distribution” model of the SEDFIT ver. 17.5 software [41], with a resolution of 300 and a confidence level (F-ratio) set to 0.68, in the range of 0.2-15.0 S. The following fitting parameters were used: Marquardt-Levenberg type algorithm and Maximum Entropy regularization. The results were plotted using the GUSSI graphical program [42].

### Determining the optimum parameters for LOca-1 peptidase activity

To determine the optimal conditions for LOca-1 peptidase activity, a series of assays were carried out under the following experimental conditions. Unless otherwise specified, a final concentration of 0.1 µg/mL of freshly activated LOca-1 was tested in activation buffer (50 mM HEPES pH 7.5, 75 mM NaCl, 1 mM CaCl_2_) at 50 °C against the fluorogenic substrate, 50 mM Boc-VPR-amc, in triplicate. Negative controls containing buffer without enzyme were included for relevant conditions to correct for background fluorescence. The assay plate was incubated, and fluorescence was measured using an EnSpire™ 2300 Multilabel Reader. Data were analysed in GraphPad Prism, as described above.

#### Temperature

The optimum temperature for LOca-1 activity was determined using a peptidase assay against Boc-VPR-amc over a range of temperatures. The tested temperatures were 0, 30, 40, 50, 55, 60, 65, 70, 80, and 100 °C. The 0 °C sample was kept on ice, and the others were incubated in a thermal cycler. The assay buffer was identical to the activation buffer. Once assembled, the reactions were incubated for 7.5 minutes at each temperature, and endpoint fluorescence was immediately measured in the plate reader. The experiment duration was informed by the initial calcium titration/activation experiment timings. To avoid errors due to sample handling time prior to measurement, the data points were run in three batches (0 to 50, 55 to 65, and 70 to 100 °C). Negative controls containing buffer without enzyme were included for each temperature point to correct for background fluorescence. Measured fluorescence values (RFU) were normalized to the highest observed value to yield relative activity (% Activity). Data were plotted as % Activity versus Temperature (°C) using GraphPad Prism.

#### Buffer system and pH

To determine the optimal pH and buffer system for LOca-1 activity, peptidase assays were performed at 50 °C using 50 mM final concentrations of four buffer systems spanning pH 4.0- 9.0: sodium acetate (pH 4.0–5.0), MES (pH 5.5–6.5), HEPES (pH 7.0–8.0), and Tris-HCl (pH 7.5–9.0). All buffers also contained 75 mM NaCl and 1 mM CaCl_2_. The enzyme and fluorogenic peptide substrate were diluted in the respective buffers prior to assay setup. Negative controls containing buffer without enzyme were included for each condition to correct for background fluorescence. Reaction velocities (RFU/s) were calculated from the linear portion of fluorescence curves measured every 20 seconds (0–522 s) and normalized to the highest observed rate to yield relative activity (% Activity). The results were plotted as % Activity versus pH using GraphPad Prism.

#### NaCl concentration

A range of NaCl concentrations was tested to determine LOca-1 tolerance to NaCl and the optimum concentration for its activity. In the peptidase assay against Boc-VPR-amc, the enzyme was assayed in activation buffer containing 2 mM CaCl_2_ and 0, 10, 50, 75, 100, 250, and 500 mM NaCl. Reaction velocities (RFU/s) were calculated from the linear portion of the fluorescence curves (0–400 s) and normalized to the highest observed rate to yield relative activity (% Activity). The results were plotted as % Activity versus log[NaCl] using GraphPad Prism.

#### Ca^2+^ concentration

A range of CaCl_2_ concentrations was tested to determine LOca-1’s dependence on Ca ^2+^ ions after activation and to identify the optimum concentration for its activity. In the peptidase assay against Boc-VPR-amc, the enzyme was assayed in activation buffer containing 0, 0.1, 0.5, 1, 2, 4, 10, and 20 mM CaCl_2_. In addition, two control samples were included to assess the effect of chelating calcium added during the activation phase. The first contained 5 mM EDTA with no CaCl_2_, and the second contained 5 mM EDTA with 1 mM CaCl_2_. The assays were run at 50 °C, and the reaction velocities (RFU/s) were calculated from the linear portion of the fluorescence curves (600–1200 s) and normalized to the highest observed rate to yield relative activity (% Activity). The results were plotted as % Activity versus each sample in GraphPad Prism.

### Peptide substrate screen

A panel of 90 fluorogenic substrates (S2 File) was prepared at 2 mM in DMSO and stored in a 384-well, low-dead-volume Echo-qualified microplate. Using an Echo Acoustic Liquid Handler (Beckman Coulter, Indianapolis, IN, United States), 200 nL of each substrate was transferred to a black 384-well low-volume assay plate (Greiner Bio-One, Monroe, NC, USA). A 1 mg/mL stock of LOca-1 was diluted to 250 µg/mL in assay buffer (50 mM HEPES, pH 8.0, 75 mM NaCl, 1 mM CaCl₂) and incubated at 60 °C for 5 min to induce activation. The activated enzyme was then diluted to 10 µg/mL in the same buffer. For each reaction, 8 µL of the diluted enzyme was added to wells containing substrate, in triplicate. Control wells received 8 µL of assay buffer without enzyme.

Plates were centrifuged at room temperature for 15 sec at 600 RPM using an Eppendorf 5810R centrifuge, then transferred to a prewarmed Synergy HTX Multi-Mode Microplate Reader (BioTek, Winooski, VT, USA) set to 50 °C. Fluorescence was monitored every 3 min and 2 sec for 2 hours using excitation/emission wavelengths of 360/460 nm. Final concentrations were 50 µM substrate and 2.5% DMSO per well. Reaction velocities were calculated from 23 consecutive fluorescence readings collected after 20- minutes. Velocities were normalized to the highest observed rate and expressed as relative activity (% Activity). The 15 peptides that yielded the highest relative activities were visualized as a bar plot in GraphPad Prism.

### Michaelis-Menten kinetics

To determine the kinetic parameters of LOca-1, initial reaction velocities were measured across a range of substrate concentrations (5.25–400 µM) using the fluorogenic substrate Boc-VPR- amc. Reactions were prepared in triplicate in a black 96-well plate at two temperature points, 50 °C and 65 °C in the “optimized” assay buffer (50 mM HEPES, pH 8, 75 mM NaCl, and 1 mM CaCl₂) with 0.1 µg/mL activated enzyme. To correct for background fluorescence, negative controls containing buffer without enzyme were included. Fluorescence was recorded every 20 seconds for 10 minutes using an EnSpire™ 2300 Multilabel Reader (PerkinElmer, Turku, Finland) with excitation at 360 nm and emission at 460 nm. Initial velocities (RFU/s) were calculated from the linear portion of each reaction curve and plotted against substrate concentration. Data were fitted to the Michaelis-Menten equation using nonlinear regression in GraphPad Prism to determine *K*_m_ and *V*_max_ values. All error values shown represent the SEM.

### Effect of arginyl-specific fluoromethylketone inhibitor

Increasing concentrations (0 to 80 μM) of the irreversible metacaspase fluoromethylketone inhibitor Z-Val-Arg-Pro-DL-Arg-fmk (AdipoGen Life Sciences, Füllinsdorf, Switzerland) were preincubated with activated LOca-1 (0.26 μg/mL) for 20 min at room temperature (Klemenčič et al., 2015). Then, Boc-QAR-amc was added to a final concentration of 50 μM, and fluorescence was measured over 1 h at 50 °C in triplicate. Negative controls containing buffer without enzyme or inhibitor were used to correct for background fluorescence. Reaction velocity was calculated and plotted as a dose–response curve, normalized to a control with no inhibitor added. The curve was fitted using GraphPad Prism.

## Results

### Discovery and phylogenetic placement of LOca-1 among metacaspase domain-containing proteins

The enzyme characterized in this study was designated LOca-1, short for “Loki’s Castle orthocaspase 1”, to reflect both its origin and classification. It was identified from co-assembled metagenomic reads derived from microbial communities inhabiting a black smoker at the Loki’s Castle hydrothermal vent field, a geothermally active site located along the Arctic Mid- Ocean Ridges. Within the metagenomic data, the gene coding for LOca-1 was found within a short 5603 bp contig, and the corresponding protein was initially annotated as a putative caspase, a 288-amino acid enzyme with an estimated molecular mass of 33.1 kDa comprising a single C14 domain according to the MEROPS database [34]. Subsequent analyses using the PFAM [43] and InterPro [44] databases identified similarity to the catalytic p20 domain of metacaspases. Together with its prokaryotic origin, these features support the classification of LOca-1 as an orthocaspase.

Amino acid sequence-based analyses indicated that proteins closely related to LOca-1 occur across diverse bacterial lineages and several archaeal taxa. Many of these homologs were recovered from aquatic habitats, including marine, hydrothermal, freshwater, and groundwater systems, but were also identified in bioreactors and plant-associated systems (Fig. 1, S3 File, S4 File). Among the analysed sequences, the closest LOca-1 homolog was a caspase-family protein encoded by *Candidatus* Zhuqueibacterota (previously known as KSB1) [45], with 100% query coverage and 90% sequence identity (GenBank: RKY84329.1). Furthermore, among characterized or enriched microorganisms, the closest relative was a caspase-family protein from *Candidatus* Methanocrinis natronophilus, an alkaliphilic methanogen capable of acetate- dependent methanogenesis, sharing 100% query coverage and 71% sequence identity (NCBI Accession: WP_316966032.1) [46].

**Fig. 1:**
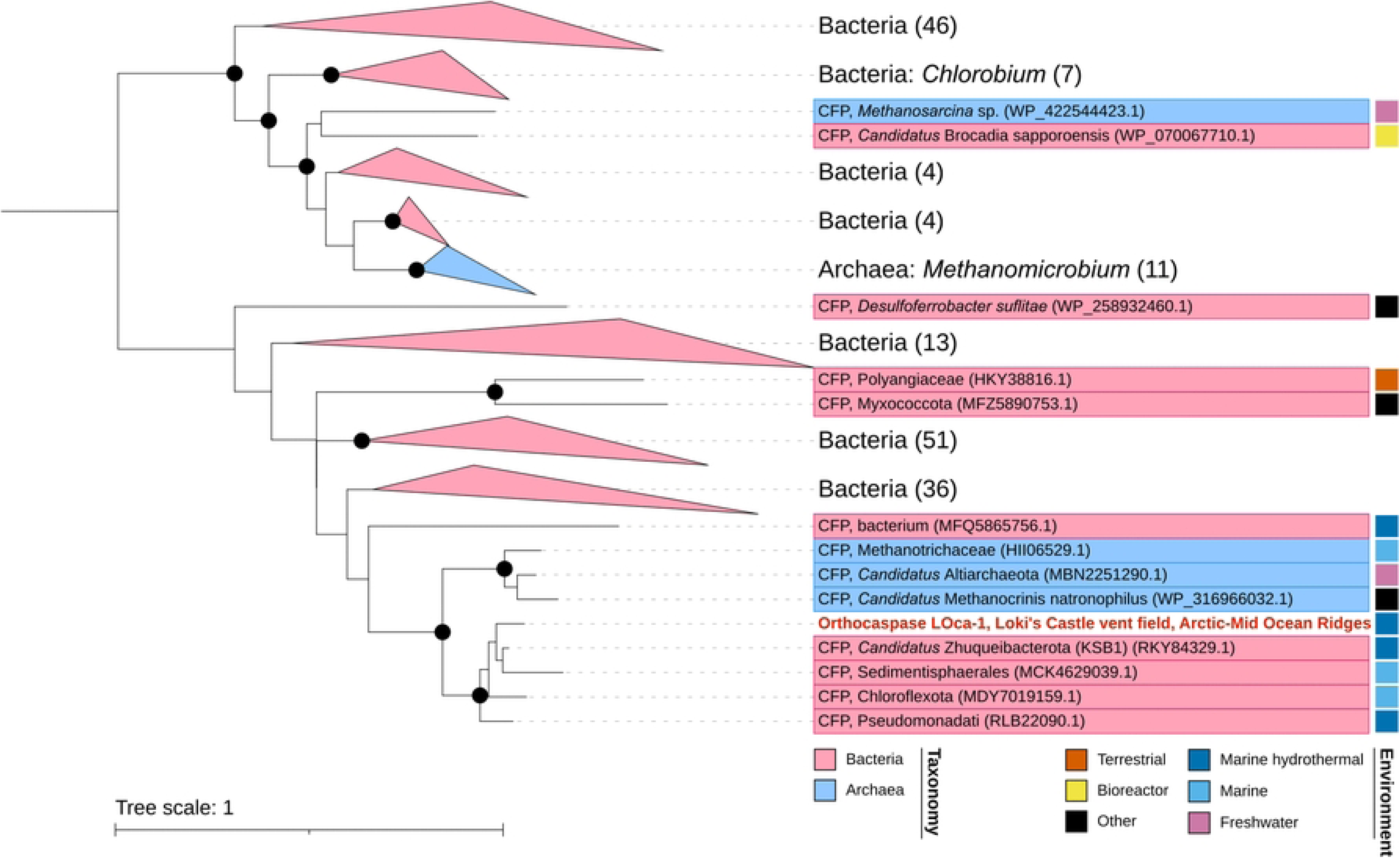
**Phylogenetic placement of orthocaspase LOca-1 from the Loki’s Castle vent field.** Maximum-likelihood phylogenetic tree of orthocaspase LOca-1 and its closest BLASTP- identified homologs. LOca-1 from this study is highlighted in red font. Taxonomic assignments and environmental metadata are based on NCBI annotations. Black circles indicate branches supported by ≥80% SH-like approximate likelihood ratio test values and ≥95% ultrafast bootstrap support (1,000 replicates each). CFP, caspase family protein.

### Structural modelling and alignment with the crystal structure of AtMCA9

Multiple sequence alignment of LOca-1 with representative metacaspases AtMCA9 from *A*. *thaliana* and AcMCA1 from *A*. *capsulatus* revealed conservation of key catalytic and regulatory motifs (Fig. 2). Despite this conservation, LOca-1 shares relatively low amino acid sequence identity with either homolog, displaying only 28% and 36.7% identity to AtMCA9 and AcMCA1, respectively. Notably, the putative activation region of LOca-1, including a conserved arginine residue corresponding to the activation site of AtMCA9, is retained in the plant metacaspase, whereas the corresponding region is absent from AcMCA1 (Fig. 2).

**Fig. 2:**
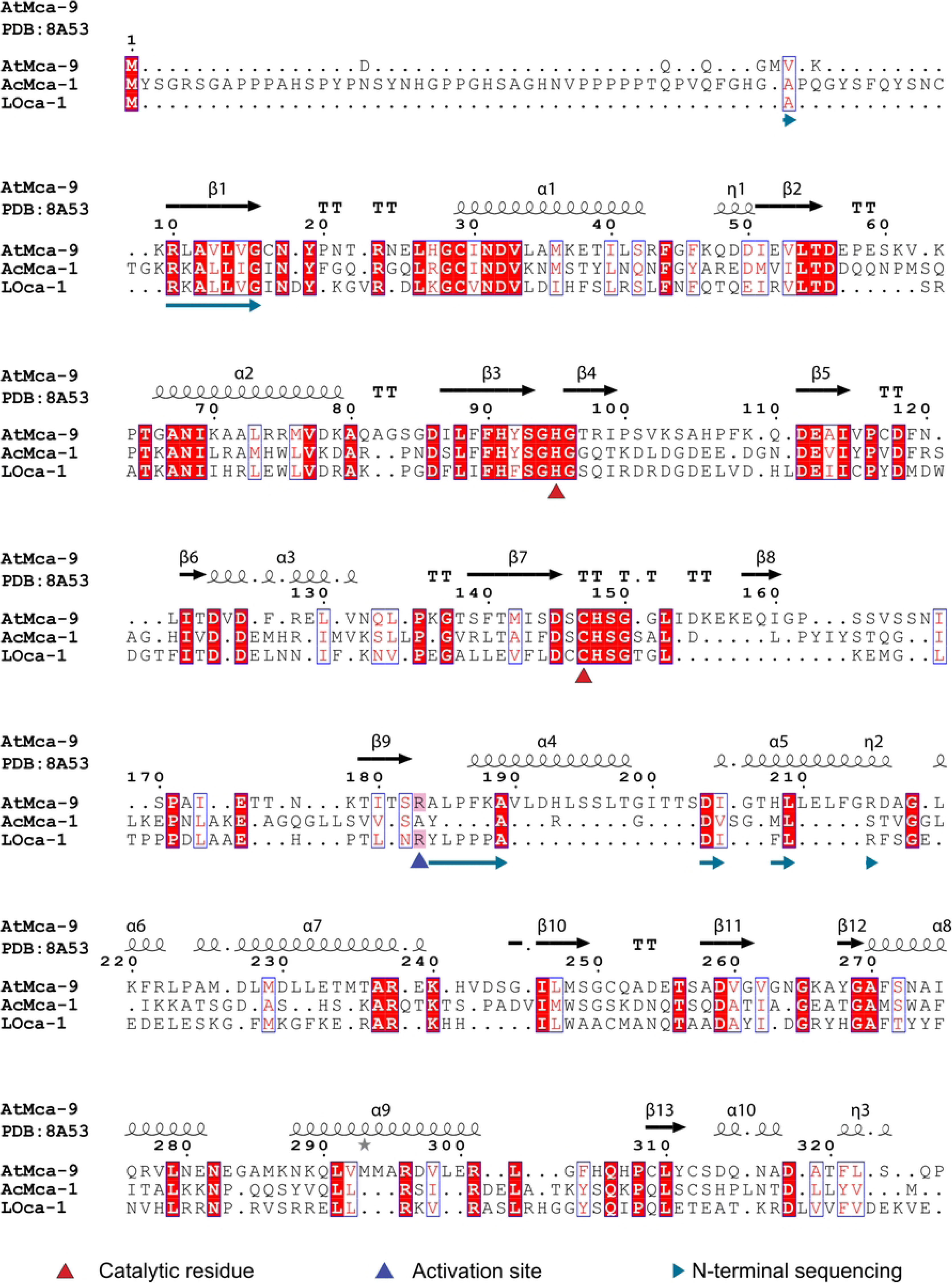
Multiple sequence alignment of LOca-1 with representative plant and fungal metacaspases from *Arabidopsis thaliana* (AtMCA9) and *Ajellomyces capsulatus* (AcMCA1). The alignment was generated using MAFFT and visualized with ESPript. Conserved catalytic residues and activation sites are highlighted. Protein numbering is based on AtMCA9 sequence and corresponds to the visualised secondary structure elements from its structure (PDB: 8A53). N-terminal sequencing of the two cleavage products generated upon LOca-1 activation confirmed proteolytic cleavage after the conserved arginine residue (Arg164). In contrast, the fungal metacaspase lacks a corresponding activation cleavage site.

A structural model of LOca-1 was generated using AlphaFold3 [36] and visualized in ChimeraX [39] (Fig. 3A). Structural alignment with the crystal structure of AtMCA9 (PDB: 8A53) showed high similarity, particularly within the catalytic core and the activation-loop region (Fig. 3B). The predicted LOca-1 structure revealed the presence of aa conserved catalytic dyad, comprising His81 and Cys138, which corresponds to the known catalytic dyad, His95 and Cys147 in *A. thaliana* (Fig. 3C). In addition, along a conserved arginine residue (Arg164) within a surface-exposed loop aligns with Arg183 in *A. thaliana* (Fig. 3C). Overall, these data support structural conservation despite the substantial evolutionary distance between bacterial orthocaspases and plant metacaspases.

**Fig. 3:**
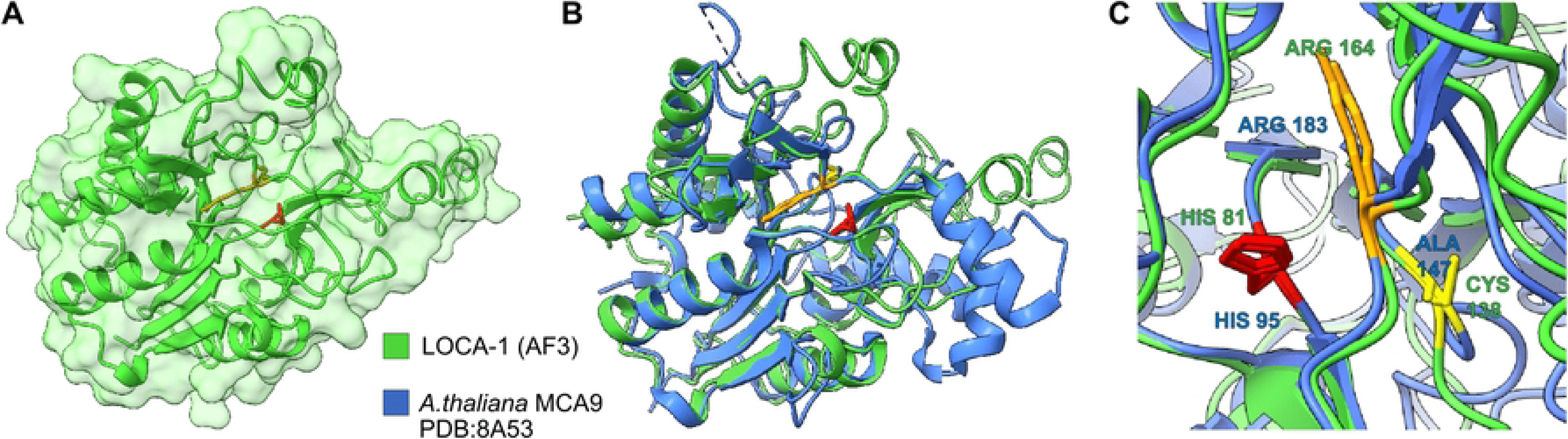
Structural model of LOca-1 and comparison with A. thaliana metacaspase 9. (A) Predicted structure of LOca-1 generated using AlphaFold3 and visualized in ChimeraX. (B) Close-up view of the catalytic dyad (histidine residues in red; cysteine and alanine residues in yellow) and the activation arginine (orange). (C) Structural superposition of LOca-1 with the crystal structure of *A. thaliana* Metacaspase 9 shows high structural similarity, particularly within the catalytic core (RMSD = 0.98 Å).

### Multimeric state and thermal stability of LOca-1

Analytical ultracentrifugation revealed a predominant sedimenting species centred at approximately 3.5 S with an estimated molecular mass of 70.4 kDa, roughly consistent with a dimeric assembly of the LOca-1 zymogen in solution. Two minor low-molecular-weight species were also detected at lower sedimentation coefficients (S5 File). The thermal stability of the LOca-1 zymogen was assessed by nanoscale differential scanning fluorimetry (nanoDSF), which revealed a single cooperative unfolding transition with an onset temperature at 66.2 ± 0.12 °C and a melting temperature (Tm) of 71.67 ± 0.03 °C (S6 Fig). Simultaneous monitoring of light scattering showed that protein aggregation was initiated at 68.51 ± 0.27 °C, and increased rapidly during thermal denaturation, reaching an inflection point at 71.87 ± 0.04 °C (S6 Fig). These results demonstrate that the LOca-1 zymogen exhibits notable thermal stability and remains structurally intact over a broad temperature range.

### Activation mechanism and activity of LOca-1

Protein activation was initially tested at pH 7.5 and 60 °C, a temperature below the measured unfolding transition of the zymogen and within the range where robust enzymatic activity was observed. Time-course SDS-PAGE analysis revealed calcium-dependent activation of LOca-1, with proteolytic cleavage products appearing as early as 2.5 min and persisting throughout the 10 min incubation period (Fig. 4A). During activation, the full-length (33.1 kDa) protein band was replaced by two shorter fragments at ∼18.5 and ∼14.5 kDa. Under similar conditions, the catalytic variant C138A did not self-activate but could be processed *in trans* upon addition of low amounts wild-type enzyme (Fig. 4B), indicating that activation is proteolytically driven. N-terminal sequencing of the two cleaved fragments reveals that the upper band corresponds to residues X to Y, while the lower band corresponded to Y to Z. This revealed that activation occurs between Arg164 and Tyr165, the same site that was predicted from the structure comparison with AtMCA9 (Fig. 3).

**Fig 4:**
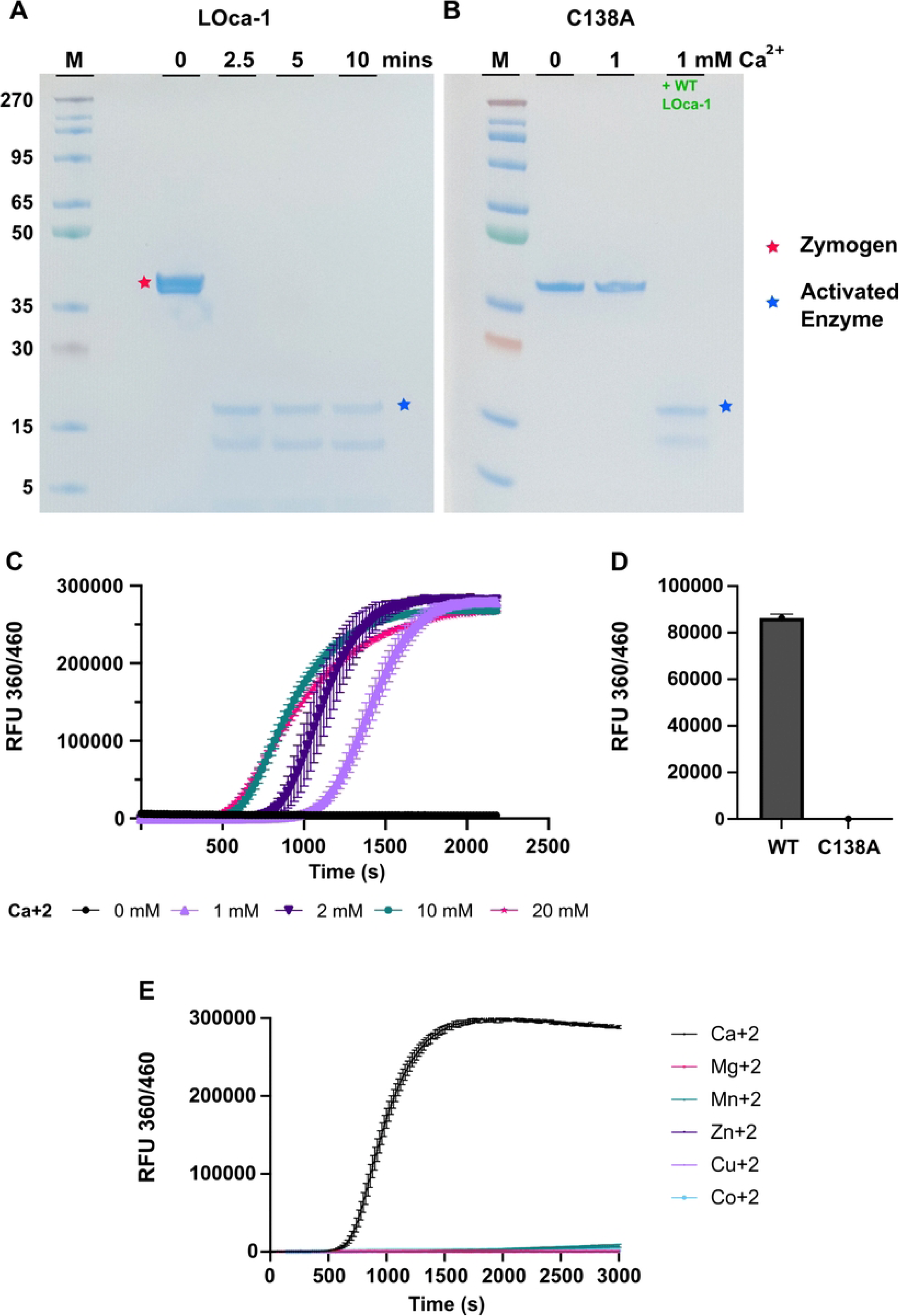
Calcium-dependent activation of LOca-1 and catalytic mutant analysis. (A) SDS–PAGE analysis showing time-dependent activation of LOca-1 at 60 °C (0, 2.5, 5, and 10 min). (B) The catalytic cysteine substitution variant C138A fails to activate autonomously in the presence of calcium but can be activated *in trans* upon addition of wild-type LOca-1. (C)

Using the fluorogenic reporter substrate Boc-VPR-amc, LOca-1 demonstrated robust peptidase activity at 60 °C only in the presence of calcium. Upon calcium titration, the time required for full enzyme activation decreased as calcium concentrations increased. The largest differences in activation timing were observed between 1 mM, 2 mM, and 10 mM CaCl₂, whereas a less pronounced change was observed between 10 mM and 20 mM CaCl₂. (Fig. 4C). Once activated, the rate of substrate cleavage was more similar in the presence of 1 to 10 mM Ca^2+^ but was slower in the presence of 20 mM. As expected, the catalytic C138A variant exhibited no detectable peptidolytic activity under identical conditions (Fig. 4D). Furthermore, LOca-1 was assayed in the presence of several common divalent metal ions to determine whether activation could also be induced by divalent cations other than calcium. Among the additional ions tested, Mg²⁺, Zn²⁺, Cu²⁺, and Co²⁺ failed to promote detectable activation. Mn²⁺ produced only a minor response relative to calcium, whereas only calcium produced rapid and robust activation (Fig. 4E). These results demonstrate a strong preference for calcium as the trigger of LOca-1 activation.

A fluorogenic substrate assay using Boc-VPR-amc demonstrates calcium-dependent activation and a time-resolved increase in fluorescence at 60 °C. (D) The C138A mutant exhibits no detectable activity compared with wild-type LOca-1 under identical conditions. All assays were performed in triplicate; error bars represent the standard error of the mean (SEM). (E) Activation and activity of LOca1 in the presence of common divalent cations confirm a strict preference for calcium.

### Optimization of LOca-1 activity conditions

To define optimal conditions for LOca-1 peptidase activity, the enzyme, upon activation, was assayed across a range of temperatures, pH values, salt and Ca ^2+^ concentrations (Fig. 5). Peak activity was observed at 65 °C, though notable activity was present even on ice (0 °C). At 70°C, 80 °C and 100°C, LOca-1 activity was reduced by approximately 2.5%, 77% and 92%, respectively, when compared to peak activity. This is consistent with the zymogen T_m_ measured at 72 °C (Fig. 5A). pH profiling showed the highest activity at pH 8.0 in HEPES and at pH 7.5 in Tris-HCl buffers, with activity sharply reduced below pH 7.5 and above 8.5. (Fig 5B). NaCl titration showed maximal activity at 75 mM (Fig 5C), but all concentrations tested (including 0 mM) yielded generally high (>75%) levels of peptidolytic activity, indicating that LOca-1 is not directly dependent on NaCl and can function well across a wide range of salinities. When a range of calcium concentrations was tested for its effect on LOca-1 activity, peak activity was observed at 1 mM CaCl₂. Even 0.1 mM of CaCl_2_ led to ∼68% of maximum activity, whereas EDTA treatment abolished activity completely, pointing to a low but necessary presence of calcium ions for LOca-1 activity (Fig. 5D). These results informed the selection of 50 mM HEPES (pH 8.0), 75 mM NaCl, and 1 mM CaCl₂ as the standard assay buffer for subsequent characterization experiments.

**Fig 5:**
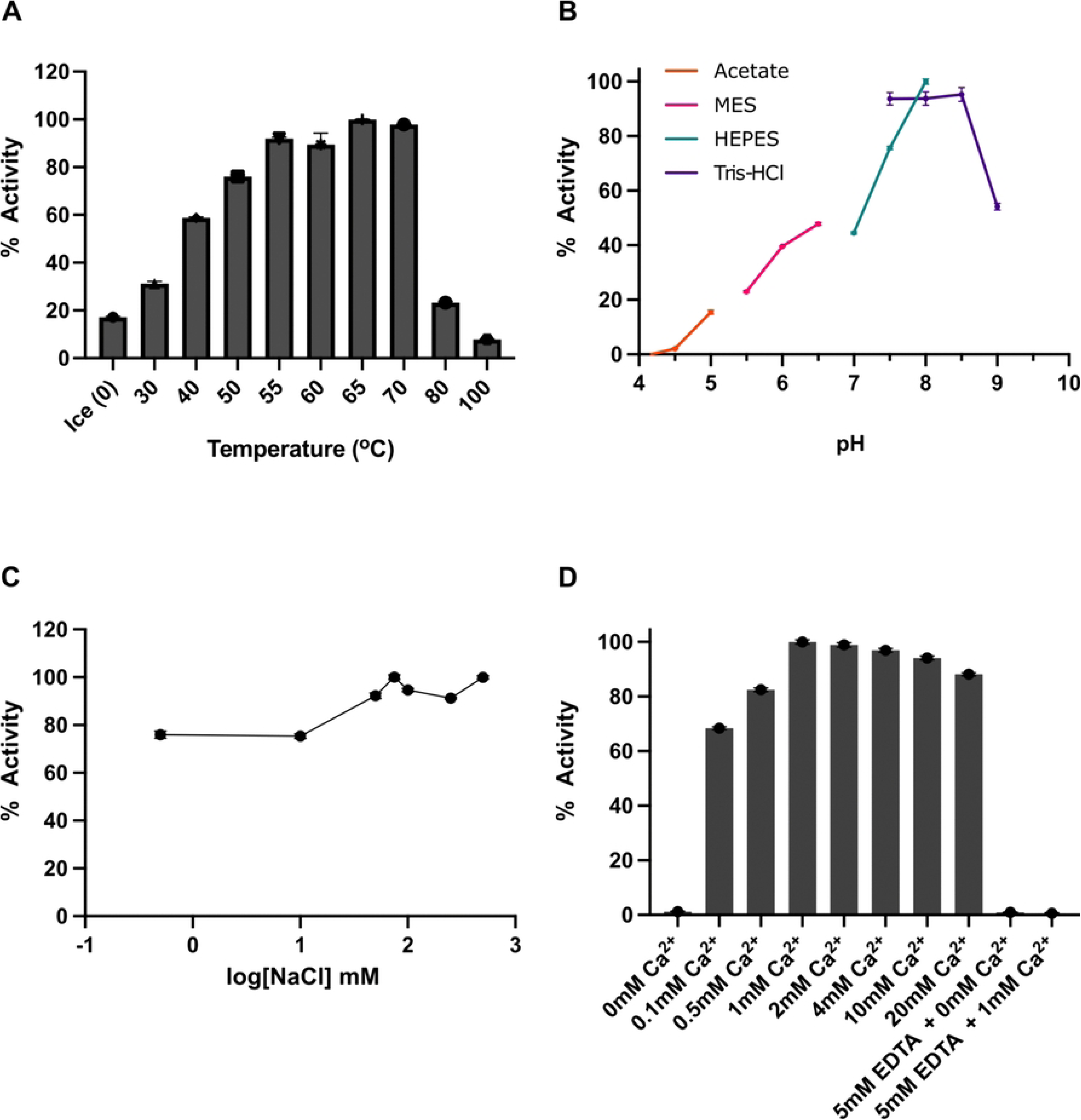
Optimization of LOca-1 activity conditions. (A) Temperature dependence of LOca-1 activity shows a broad activity range from 0 °C to 100 °C with maximal activity at 65 °C. (B) pH dependence shows optimal activity at pH 8.0 in HEPES buffer and at pH 7.5 in Tris–HCl buffer. (C) NaCl titration shows broad salt tolerance with maximal activity at 75 mM NaCl. (D) Calcium titration identifies optimal activity at 1 mM Ca²⁺, and EDTA treatment confirms calcium dependence. All assays were performed in triplicate; error bars represent the standard error of the mean (SEM).

### Substrate specificity and kinetic characterization

A broad fluorogenic peptide screen revealed that LOca-1 preferentially cleaves arginyl- containing substrates, and, surprisingly, Boc-VPR-amc yielded the highest activity among the 90 peptides tested (Fig 6A, S2 File). LOca-1 did not process any substrates on the C-terminal side of lysine. Michaelis-Menten kinetics were analysed both at 50 and 65 °C using Boc-VPR- amc and the resulting *K*_m_ values were 35.95 and 71.52 µM, respectively (Fig 6B). In addition, the *V*_max_ values were 97.55 RFU/s (50 °C) and 163.5 RFU/s (65 °C). Inhibition by the arginyl- specific fluoromethylketone inhibitor z-VRPR-fmk confirmed potent inhibition of LOca-1 activity with an IC_50_ of 19.6 nM (Fig. 6C).

**Fig 6:**
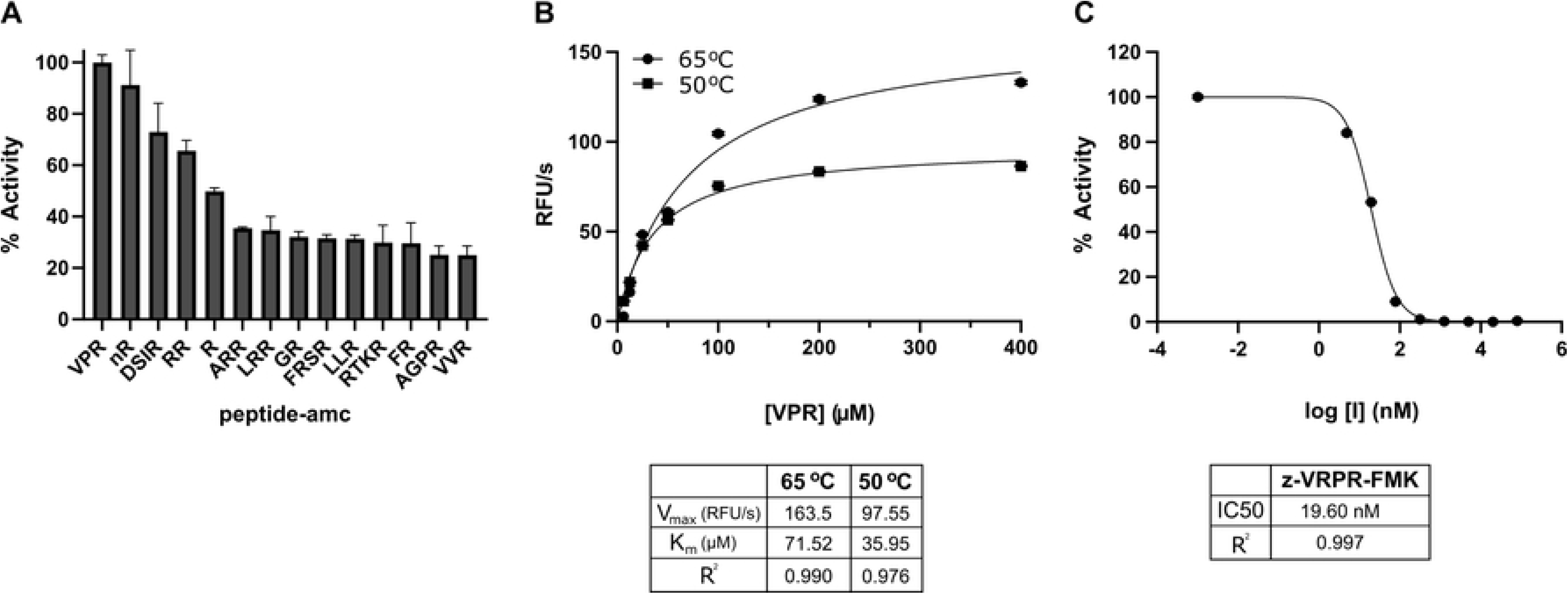
Substrate specificity and kinetic parameters of LOca-1. (A) Activity of LOca-1 against a panel of 90 fluorogenic peptide substrates shows a strong preference for arginyl-containing sequences, with Boc-VPR-amc exhibiting the highest and most consistent activity. The top fifteen substrates are shown. (B) Michaelis–Menten kinetics for Boc-VPR-amc measured at 50 °C and 65 °C; corresponding *K*◻ and *V*◻ₐₓ values are indicated. (C) Dose–response analysis of the inhibitor z-VRPR-fmk shows complete inhibition of LOca-1 activity at nanomolar concentrations. All assays were performed in triplicate; error bars represent the standard error of the mean (SEM).

## Discussion

In this study, we present the characterization of LOca-1, a minimal domain orthocaspase identified from the Loki’s Castle hydrothermal vent field. Our analyses reveal a thermostable, calcium-activated, arginyl-specific protease that expands the limited number of experimentally characterized orthocaspases. While caspases and metacaspases have been extensively characterized in eukaryotes for their roles in programmed cell death and stress responses [2,47], orthocaspases remain comparatively poorly studied. To our knowledge, only one other orthocaspase, MaOC1 from the freshwater cyanobacterium *Microcystis aeruginosa*, has been biochemically characterized and experimentally verified to possess proteolytic activity beyond bioinformatic predictions [7,12]. In this context, LOca-1 represents only the second experimentally validated orthocaspase and the first derived from a marine deep-sea hydrothermal environment, expanding the known ecological and biochemical diversity of orthocaspases. Phylogenetic analyses place LOca-1 within a diverse clade of orthocaspases recovered from environmental samples, including marine sediment-derived lineages (Fig 1). This placement is consistent with previous reports describing the broad phylogenetic and ecological distribution of orthocaspase homologs across prokaryotic taxa [8,48].

While all orthocaspases share a conserved p20 catalytic core, many exhibit considerable architectural diversity through the presence of additional regulatory, sensory, and protein interaction domains [8,9,48]. In contrast, LOca-1 consists primarily of the p20 catalytic domain, representing the simplest orthocaspase architecture currently described. This organization renders LOca-1 a particularly valuable model for defining the core biochemical properties of orthocaspases and establishing a benchmark against which the properties of other orthocaspases and the regulatory contributions of relevant accessory domains can be assessed.

In this work, we demonstrated that LOca-1 undergoes rapid, calcium-dependent activation through cleavage after a conserved arginine residue (Arg164) at 60 °C. However, elevated temperatures alone were insufficient to activate LOca-1 in the absence of calcium, showing that calcium is essential for enzyme maturation (Fig 4). Furthermore, the strong preference for calcium over other divalent cations signifies a specific role for calcium in LOca-1 activation, rather than a general requirement for metal ions. Calcium-dependent regulation is recognized as an important feature of prokaryotic physiology, with transient changes in intracellular Ca²⁺ concentrations associated with diverse cellular processes, including chemotaxis, motility, stress responses, sporulation, biofilm formation, and cell cycle regulation [49]. Despite the clear dependence of LOca-1 on calcium for activation, sequence-based bioinformatic analyses detected no canonical calcium-binding motifs or architectures, including EF-hand domains [49,50]. Beyond its role in autocatalytic maturation, calcium had only a limited influence on enzymatic activity. Relatively low calcium concentrations were sufficient to support maximal proteolytic activity, which did not increase further at higher calcium concentrations. The molecular basis of calcium sensing in LOca-1 nevertheless remains unresolved, as calcium- responsive proteins may utilize non-canonical binding sites or distributed structural features that are not readily predicted from primary sequence alone [49].

On a broader scale, this calcium-dependent activation mechanism is shared with notable fungal [51,52] and plant metacaspases, including AtMCA4, AtMCA5, and AtMCA8 from *Arabidopsis thaliana* [53–55]. Structural comparison of the LOca-1 AlphaFold3 model with the crystal structure of AtMCA9 (PDB ID: 8A53)[38] displayed notable similarity in both the catalytic architecture and activation-loop region despite the substantial evolutionary distance separating bacterial orthocaspases and plant metacaspases. Notably, despite sharing a similar p20 catalytic core, the two enzymes employ distinct regulatory triggers, with AtMCA9 being activated through a pH-dependent mechanism and LOca-1 through calcium-dependent proteolytic maturation [56].

In comparison, the only previously characterized orthocaspase, MaOC1, undergoes calcium- independent autocatalytic activation and is a substantially larger protein, with an extended C- terminal region beyond the conserved p20 catalytic core [7,12]. The contrasting activation mechanisms of LOca-1 and MaOC1 demonstrate that orthocaspases can employ distinct regulatory strategies despite sharing a common p20 catalytic scaffold. Moreover, the marked differences in overall protein architecture between the two enzymes suggest that structural features beyond the catalytic core may contribute to differences in activation mechanisms. Together with the distinct pH-dependent activation described for AtMCA9, these observations emphasize that diverse members of the metacaspase family can retain a conserved catalytic architecture while employing different regulatory triggers [2,56].

Analytical ultracentrifugation further revealed that the LOca-1 zymogen primarily exists as a dimer in solution. To our knowledge, this finding represents the first experimental evidence for the oligomeric state of an orthocaspase, and for the conserved p20 catalytic domain itself. This contrasts with characterized metacaspases, which are generally described as monomeric enzymes whose activation relies on calcium-dependent conformational changes and proteolytic maturation rather than oligomerization [51,52,55]. Unlike initiator caspases, for which oligomerization is central to activation [57], the dimeric LOca-1 zymogen remained entirely dependent on calcium-mediated proteolytic maturation. Together, these observations suggest that oligomeric organization of the LOca-1 zymogen alone is insufficient to enable autoproteolytic activation. Future structural characterization of both the zymogen and activated forms of LOca-1 may provide important mechanistic insight into the structural basis of calcium- dependent activation and how proteolytic cleavage at Arg164 reorganizes the catalytic machinery into its active state.

LOca-1 shows strict specificity for arginine at the P1 position, with no detectable activity against lysine-containing substrates despite screening a broad fluorogenic peptide panel. Boc-VPR-amc was identified as the preferred substrate, and the arginyl-specific fluoromethylketone inhibitor z-VRPR-fmk potently inhibited enzymatic activity at nanomolar concentrations. While these findings closely align with those reported for MaOC1 [7], LOca-1 and MaOC1 also differ in their relative preferences for residues preceding the scissile arginine, suggesting that although P1 specificity is conserved, substrate recognition beyond the P1 position can vary among orthocaspases. In LOca-1, residues at the P2 position appear to play a major role in determining cleavage efficiency, consistent with observations for other cysteine proteases such as papain and globupain [40,58]. Together, these observations indicate that while fine-scale substrate preferences may vary among orthocaspases, recognition of arginine- containing substrates remains a conserved functional trait of the p20 catalytic domain. Although LOca-1 exhibits a particularly strict requirement for arginine, this specificity remains consistent with the broader tendency of metacaspase-family proteases to recognize basic residues [2].

LOca-1 possesses notable thermal stability, retaining activity across a broad temperature range and showing maximal activity at 65 °C, with a melting temperature of approximately 72 °C. Although the biochemical properties of LOca-1 broadly parallel those of other characterized metacaspases and orthocaspases, its thermal profile distinguishes it from the only previously characterized orthocaspase, MaOC1[7,12]. Its ability to function under both mesophilic and thermophilic conditions suggests that substantial thermal adaptation can occur without compromising the defining catalytic properties of the p20 domain, highlighting the functional flexibility of an otherwise conserved catalytic module.

LOca-1 originates from the Loki’s Castle hydrothermal vent field, an environment characterized by steep physicochemical gradients and dynamic mixing of hydrothermal fluids with seawater [59]. Microorganisms and their enzymes often exhibit adaptations that promote stability and function under prevailing physicochemical conditions [15–17]. The combination of broad thermal tolerance and calcium-dependent activation observed in LOca-1 may therefore reflect an adaptation to its native environment. Because hydrothermal fluids at Lokís Castle are characterized by both elevated temperatures and higher calcium concentrations than surrounding seawater [60], the calcium-dependent activation of LOca-1 is particularly noteworthy. However, whether calcium-dependent activation represents an adaptation to calcium-rich hydrothermal environments or reflects a more ancient regulatory property of orthocaspases remains unclear. Broader functional characterization of orthocaspases across phylogenetically distinct taxa and contrasting environments may provide new insight into how protease activation strategies have evolved and diversified throughout the family.

Although the precise physiological factors governing LOca-1 activation, substrate recognition, and biological function remain unknown, the combination of regulated enzyme maturation and highly selective substrate recognition is more consistent with controlled proteolysis than with general protein turnover [2,12]. Across plants, fungi, and protists, members of the metacaspase family have been implicated in diverse regulatory processes including programmed cell death, stress adaptation, developmental transitions, and cell cycle-associated pathways [2,3,55]. Even though direct functional evidence remains scarce for prokaryotic homologs, the orthocaspase MaOC1 has similarly been proposed to participate in regulatory networks associated with a toxin-antitoxin system [12]. Together, these observations support the hypothesis that orthocaspases participate in regulated proteolytic processes rather than general protein turnover.

An important unresolved question is whether orthocaspases have a conserved ancestral biological function across lineages, or whether the p20 protease domain functions as a versatile catalytic module that can be adapted for participation in distinct regulatory networks [2,9]. Likewise, how secondary domains beyond the catalytic core influence the observed diversity of molecular regulation and associated cellular functions remains unclear. Addressing these questions will require further biochemical, genetic, and comparative genomic investigation of LOca-1, together with broader characterization of orthocaspases from diverse microbial lineages.

## Conclusion

In this work, we identify and characterize LOca-1, a novel orthocaspase from the Lokís Castle hydrothermal vent field. Our results reveal a biochemical profile defined by calcium-dependent activation, strict arginyl specificity, and substantial thermal stability. As one of only two experimentally characterized orthocaspases, LOca-1 expands the known biochemical diversity of prokaryotic metacaspase homologs while providing insight into the properties of a minimal p20-domain orthocaspase. These findings establish a foundation for future studies aimed at identifying physiological substrates, clarifying biological function, and understanding the roles of orthocaspases across diverse microbial lineages.

## Acknowledgements

This work was funded by the InDepth project through the Norway Financial Mechanism and the National Science Center (Poland) GRIEG1 grant: UMO-2019/34/H/NZ2/00584, the Research Council of Norway through the DeepSeaQuence (project number 315427), and has also received support through the Trond Mohn Foundation, the University of Bergen through the Centre for Deep Sea Research (grant #TMS2020TMT13). We thank Håkon Dahle from the Computational Biology Unit at the University of Bergen for the original metagenomic assembly that contained LOca-1.

## Supporting information

S1 File: Fasta file containing the DNA and amino acid sequences of LOCa-1.

**S2 File: Spreadsheet displaying the full peptide substrate screening panel.** Activity results for each tested peptide are sorted according to the mean reaction velocities calculated from 23 consecutive fluorescence readings collected between 20- and 100-minute time points.

**S3 File: Phylogenetic placement of orthocaspase LOca-1– full-size figure.** Phylogenetic tree of orthocaspase LOca-1 including closest relative reference proteins using BLASTP. LOca-1 from this study is shown in red. The taxonomy and environments specified are based on NCBI metadata. Black node circles mark branches with support values higher than 80% with SH-like approximate likelihood ratio test and 95% with ultrafast bootstrapping, both including 1000 iterations.

S4 File: Spreadsheet containing the relevant metadata of each entry in Fig 1 and S3 File.

**S5 File: Sedimentation velocity analytical ultracentrifugation of the LOca-1 zymogen.** Continuous c(s) distribution analysis revealed a predominant species at approximately 3.5 S corresponding to an estimated molecular mass of 70.4 kDa, roughly consistent with a dimeric assembly of LOca-1 in solution.

**S6 Fig: NanoDSF profiles of LOca-1.** The thermal unfolding profile of LOca-1 zymogen as determined by nanoDSF. The enzyme exhibited a single cooperative unfolding transition with an unfolding onset at 66.2 ± 0.12 °C and a melting temperature (T_m_) of 71.67 ± 0.03 °C. The onset of aggregation (Tagg) was observed at 68.51 ± 0.27 °C.

